# Systematic detection of amino acid substitutions in proteome reveals a mechanistic basis of ribosome errors

**DOI:** 10.1101/255943

**Authors:** Ernest Mordret, Avia Yehonadav, Georgina D Barnabas, Jürgen Cox, Orna Dahan, Tamar Geiger, Ariel B Lindner, Yitzhak Pilpel

## Abstract

Translation errors limit the accuracy of information transmission from DNA to proteins. Selective pressures shape the way cells produce their proteins: the translation machinery and the mRNA sequences it decodes co-evolved to ensure that translation proceeds fast and accurately in a wide range of environmental conditions. Our understanding of the causes of amino acid misincorporations and of their effect on the evolution of protein sequences is largely hindered by the lack of experimental methods to observe errors at the full proteome level. Here, we systematically detect and quantify errors in entire proteomes from mass spectrometry data. Following HPLC MS-MS data acquisition, we identify *E. coli* and *S. cerevisiae* peptides whose mass and fragment ion spectrum are consistent with that of a peptide bearing a single amino acid substitution, and verify that such spectrum cannot result from a post-translational modification. Our analyses confirm that most substitutions occur due to codon-to-anticodon mispairing within the ribosome. Patterns of errors due to mispairing were similar in bacteria and yeast, suggesting that the error spectrum is chemically constrained. Treating *E. coli* cells with a drug known to affect ribosomal proofreading increased the error rates due to mispairing at the wobble codon position. Starving bacteria for serine resulted in specific patterns of substitutions reflecting the amino acid deficiency. Overall, translation errors tend to occur at positions that are less evolutionarily conserved, and that minimally affect protein energetic stability, indicating that they are selected against. Genome wide ribosome density data suggest that errors occur at sites where ribosome velocity is relatively high, supporting the notion of a trade-off between speed and accuracy as predicted by proofreading theories. Together our results reveal a mechanistic basis for ribosome errors in translation.

## Introduction

Genetic information propagation along the central dogma is subject to errors in DNA replication, RNA transcription and protein translation. DNA replication typically manifests the highest fidelity among these processes, featuring genetic mutation rate on the order of 10^−9^ - 10^−10^ per nucleotide per genome doubling^1,2^. “Phenotypic mutations”, i.e. errors in RNA transcription and protein translation, in which the wrong RNA nucleotide or amino acid are respectively incorporated, occur at considerably higher rate. Bacterial RNA polymerases misincorporate nucleotides every 10^4^ to 10^5^ positions *in vivo*^3^. In *E. coli*, amino acid substitutions rates were precisely measured at defined sites using a luciferase reporter construct^4^, to typically reside within the 10^−4^ – 10^−3^ per position range.

Since the translation machinery relies on tRNAs to match amino acids with codons, errors can result either from the charging of a tRNA with a non cognate amino acid (synthetase error, or “mischarging”), or from the ribosome failing to discriminate against imperfect codon-anticodon complexes in its A-site (ribosome error, or “mispairing”). The accuracy of both processes is amplified by kinetic proofreading^5,6^, a general mechanism during which the addition of an irreversible, energy consuming step to a reaction permits the system to reach discrimination levels that are inaccessible at thermodynamic equilibrium. Theoretical models demonstrated that processes relying on kinetic proofreading to increase their accuracy were necessarily subject to a trade-off between incorporation speed, accuracy and energetic cost^7–9^.

The amount of resources that cells invest to ensure that proteins function properly indicates that errors during protein synthesis can strongly affect fitness. Indeed, proteins that contain amino acid substitutions tend to misfold and aggregate, promote spurious protein-protein interactions, and saturate the protein quality control machinery, resulting in proteotoxic stress^10^, and potentially in diseases^11^. Conversely, some errors might even prove to be advantageous: moderate levels of methionine misacylation on various non-methionine-tRNAs were shown to provide a fitness advantage in response to oxidative stress from bacteria to humans^12–14^. As example, high error levels across instances of rare codon allow a parasitic yeast to increase its adherence and its ability to evade the immune response^15^. It this case, mistranslation is beneficial in response to environmental stresses as it can help sustain and disseminate cellular phenotypic viability, e.g. as it may enhance pathogens survival by creating antigenic diversity in surface proteins^15^ On an evolutionary time scales too, phenotypic errors might open evolutionary paths otherwise precluded by epistatic interactions^16^, and facilitate the purging of deleterious mutations^17^. A computational analysis of codon usage patterns across genomes revealed that natural selection constrains the identity of codons at evolutionarily conserved positions, suggesting that evolution favors more accurate codons at these sites^18^.

Whereas the rates of DNA mutation and RNA polymerase errors are now quantifiable across the genome and the transcriptome thanks to next-generation sequencing, errors in protein translation have remained elusive. An early effort by Edelmann and Gallant^19^, who quantitatively tracked the insertion of radioactively labeled cysteine in *E. coli*’s flagellin, a cysteine free protein, revealed a first global estimate of mistranslation, with misincorporations happening on average every 10,000 amino acids. Since then, the use of luminescent reporter constructs allowed the quantitative tracking of specific types of mistranslation, at defined positions within these probes^4,20^. These methods have highlighted the importance of codon-anticodon recognition and tRNA competition as determinants of these error rates, and were used to characterize the effects of aminoglycoside antibiotics and ribosome ambiguity mutations (ram)^4,20^.

However, major questions still remain open. While error rates could be quantified at specific sites, the overall error spectrum across the proteome has not yet been characterized. Such measurements would shed light on the relative contribution of aminoacyl tRNA Synthetase (aaRS) and ribosomes to translation errors. Furthermore, detecting and quantifying amino acid substitutions across the proteome may reveal evolutionary constraints imposed by translation infidelity and allow us to ask how organisms locally modulate their error levels, and the strategies they employ to mitigate their deleterious effects.

Mass Spectrometry (MS)-based proteomics, enables routine, high throughput characterization of canonical proteomes and common post translational modifications (PTMs), and was described as an upcoming tool for the study of protein mistranslation for almost a decade^10^. Recently, proteomics was harnessed to detect various substitutions from several purified recombinant proteins ^21^ and to track the incorporation of norvaline at leucine positions across the proteome of E. coli mutants^22^. Yet, MS has yet to be harnessed for the systematic study of amino acid substitutions on a proteome wide scale. Such study was hitherto hindered by the low abundance of substitutions compared to other natural and post-translational protein modifications, and a much larger search space.

To identify these rare translation errors, we performed a deep proteomic analysis of *E. coli* cells, and assessed the effects of an aminoglycoside antibiotic (paromomycin) on the bacteria’s translation error rates and spectrum. In addition, we also tested the effect of serine starvation on the mistranslation rate of its cognate codons. We carried out our analysis with MaxQuant^23^, repurposing its dependent peptide algorithm to identify mass shifts consistent with amino acid substitutions, and stringently filtering out potential methodology artifacts. We then validated these identifications using a set of independent analyses that include a shift in HPLC retention time due to change in hydrophobicity of the encoded amino acid. Performing these experiments and analyses on *E. coli* in several growth conditions and analyzing similar data in *S. cerevisiae* ^24^, we could detect over 3,500 independent substitution events.

This dataset, unprecedented in its type and extent, allowed us to begin to unravel the mechanistic basis of translation errors. We found that most errors result from mispairing between codons and near-cognate tRNAs, mostly within the A site of the ribosome. We could derive the amino acid error spectrum of each codon in the genetic code, and deduce patterns of mispairing between codons and anticodons at each of the three codon positions. We could also measure the effect of the aminoglycoside on the error pattern and revealed that it mainly causes errors due to mispairing at the third codon position. We found that error spectra in yeast and bacteria are similar, probably reflecting shared chemical constraints. We found that errors are allowed to occur more frequently at evolutionarily rapidly evolving sites, at protein structure sites in which mutations are more tolerated energetically, and in positions in which the ribosome progresses more rapidly. Our results demonstrate that natural selection has tuned translation error rates at specific protein positions, acting upon the nucleotide sequence of proteins to reduce the cost burden of translation errors when most detrimental.

## Results

### A pipeline to confidently identify amino acid substitutions in a proteome

Mass spectrometry enables the fast, large-scale identification of peptides within complex samples. From a computational point of view, amino acid substitutions can be regarded as a particular case of post-translational modifications (PTMs), many of which are now routinely studied at the proteome level. However, the standard database search algorithm is not adapted to the large scale detection of substitutions: assuming peptides of average length of 10 amino acids, there would be on the order of 200 times more singly-modified than canonical peptides to search for, leading to impractical search times and a considerable loss of statistical power.

Blind modification searches^25–27^ offer a way to identify modified peptides without requiring the user to input a list of predefined modifications. They rely on the observation that modified peptides are usually less abundant than their unmodified counterparts, and are therefore only likely to be present in the sample if the unmodified peptide has already been detected. We thus adapted MaxQuant^23^, developed previously for the analysis of post-translational modifications in order to identify mistranslated peptides using its “dependent peptide search” algorithm; see outline of our pipeline in Figure 1A. “Dependent Peptides” are defined as peptides that show mass shifts in comparison to the unmodified, genome-encoded “Base Peptides” (Fig. 1B). We then applied a series of filters to the list of dependent peptides, in order to stringently remove known PTMs and known artifacts and conservatively retain only amino acid substitutions. For a detailed description of the pipeline, see Methods.

**Figure 1:**
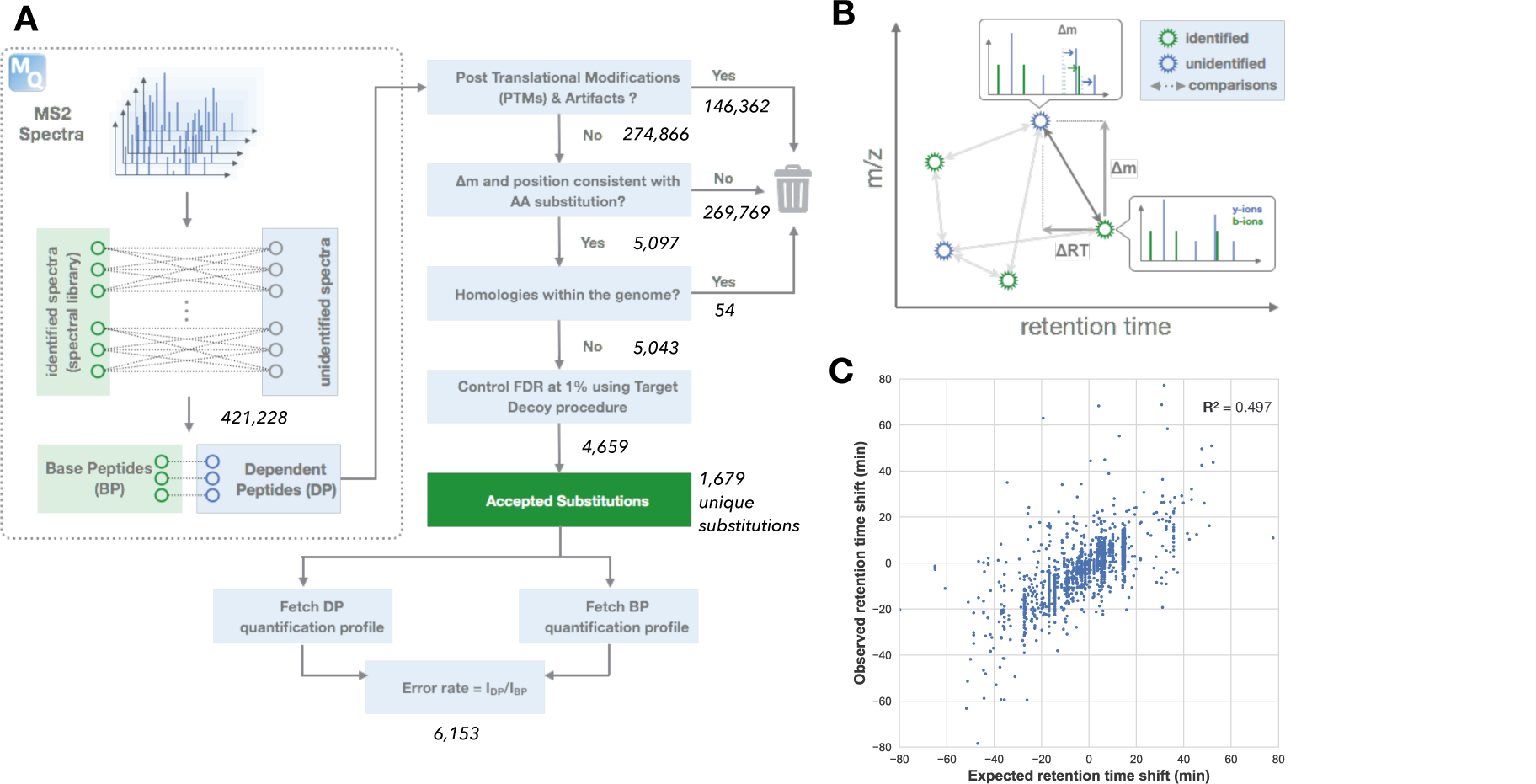
a computational pipeline to confidently identify amino acid substitutions from Mass Spectrometry data. A: Overview of the pipeline. For a detailed description of the different steps, see Material & Methods. B: MaxQuant Dependent Peptide search performs exhaustive pairing of unidentified spectra to a spectral library derived from the identified spectra. For each pair of (identified, unidentified) spectra of the same charge z, and found in the same fraction, the algorithm first computes the mass difference Δm = m_unidentified_ - m_identified_. It simulates *in silico*, and sequentially, the addition of a single moiety of mass Δm at any position in the identified peptide, and generates the corresponding theoretical spectrum for the modified peptide. These spectra are then compared to the experimental spectrum using MaxQuant Andromeda’s score formula. The pair with the highest score is retained, and the significance of the match is assessed using a target-decoy FDR procedure. C: The observed retention time shift induced by our set of substitutions is accurately predicted by a simple sequence-based retention time model.

The detection of translation errors, rare events by definition, required specific experimental settings. Mass spectrometers sample in priority the most abundant peptides for MS2 fragmentation. Since we expect translation errors to be present at much lower level than error-free peptides, we fractionated our samples to reduce their complexity and increase the chances of sampling low abundance peptides. We separated proteins into a high solubility and a low solubility fraction^28^, which we further fractionated using strong cation exchange (SCX) chromatography.

We generated two deep-coverage, high-resolution maps of the *E. coli* proteome. In our first dataset, we measured the effects of paromomycin at sub-lethal concentration on the ribosome’s accuracy, in rich conditions (LB, 37°C), and compared it to a non-treated sample. In the second dataset, we starved a serine auxotroph for serine (MOPS, 37°C) and compared its substitution pattern to the parental unstarved wild type. In total we generated error maps of nine samples, each in two replicates (see Methods). Altogether we detected 3,596 independent amino acid substitutions (each defined here by a unique position within a specific protein and a unique amino acid substitution destination)) in the *E. coli* proteome. In addition to the sixteen samples from the bacterium we also analyzed an existing proteome dataset^24^ from the yeast *S. cerevisiae* at a single type, non-treated, condition that yielded 225 independent substitutions.

### Most of the high quality hits are bona fide amino acid substitutions

First, we grew wild type *E. coli* cells (MG1655) in defined medium (MOPS complete, 37°C) in biological duplicates, harvested cells at three time points, roughly corresponding to early exponential phase, late exponential phase, and stationary phase, and used our pipeline to detect amino acid substitutions. Given mass differences detected between base and dependent peptides we must first establish that they represent *bonafide* amino acid substitutions, and not methodological artifacts. We took advantage of the fact that many amino acid substitutions result in a change of peptide hydrophobicity and they should hence result in retention time shifts during liquid chromatography. The retention time of a peptide can be predicted with high accuracy (R^2^ > 0.9) as the sum of the hydrophobicity coefficients of its amino acids^29^. Therefore, the predicted HPLC retention time of the substituted amino acid can be computed and compared to the observed retention time recorded for the substituted peptide. We trained a retention time prediction tool^30^ on a list of confidently identified unmodified peptides, and used it to generate an expectation of the retention time shift induced by the substitutions. We compared this expectation to the observed retention time shift for each of the detected substitutions (Fig. 1C). We observed a good agreement between the observed retention time shifts and our model, supporting the notion that most of the substitutions detected are genuine amino acid replacements.

Note that, since MS2 spectra are systematically recorded for highly abundant parent ions, our sampling strategy is biased towards the detection of substitutions originating from highly expressed proteins.

We define a substitution as a combination of a position in a protein, characterized by an “origin” amino acid (and its associated codon), and a “destination” amino acid. We then divide all substitutions in two sets: a substitution is classified as a Near Cognate Error (NeCE) if the error-bearing codon of the origin amino acid matches two out of three bases of one of the codons of the destination amino acid, and as Non Cognate Error (NoCE) if more mismatches would be required. For example, a substitution from Serine encoded by AGC into an Asparagine is considered NeCE since of the codons encoding this amino acid destination AAC, represents a single mismatch (at the 2^nd^ position); in contrast substitution of a Cysteine encoded by a UGU into an Alanine must be a NoCE since none of the Alanine codons (GCN) matches with a single mismatch. The structure of the genetic code dictates that only a minority of the substitutions would be classified as NeCE. In particular, of all detectable codon to amino acid substitution types 30% are expected to be of the NeCE type. In stark contrast, 88% of the unique substitutions detected by our method with the full *E. coli* dataset are classified as NeCE. Thus, the great majority of observed substitutions in our data can be rationalized by a similarity between the origin’s codon and a codon of the destination amino acid. Such enrichment for NeCE serves as an indication that we inspect genuine amino acid substitutions.

The ribosome uses small differences in free energy between correct and incorrect codon-anticodon matches to select tRNAs, and was shown to generate NeCE at much higher rates than NoCE^4^. During loading of an amino acid by an aaRS, an error can stem from the choice of an incorrect tRNA or the loading of an incorrect amino acid. Since the majority of synthetases assess the identity of the tRNA by probing the bases of the anticodon, this first mechanism of error is also likely to generate mostly NeCE. However, if the error results from the loading of the wrong amino acid, there should be a priori no preference for NeCE over NoCE. We will consider NoCE as more likely to stem from a synthetase error, and examine the notion that the majority of NeCE indeed represent mRNA-tRNA mispairing events.

### Overview of E. coli’s amino acid substitution landscape

We generated 64*20 codon to amino acid matrices that depict the prevalence of each type of amino acid error in a dataset. Note that these matrices do not represent real relative probabilities of errors as they are affected by biases such as codon usage. The numbers of unique peptides supporting any codon to amino acid substitution type is presented in Fig. 2A; the intensity of the shade is proportional to the logarithm (base 2) of the number of unique genome positions in which a given substitution type was observed. Because leucine and isoleucine are isomers and thus share the exact same mass, our method is not able to distinguish the two amino acids as destinations of a substitution, and we treat them as a single destination amino acid. Furthermore, substitution types that transform a codon into its cognate amino acid, involve a stop codon, or substitutions that cannot be detected using our method because they represent a mass shift that corresponds precisely to the mass shift and specificity of at least one known PTM, were grayed out, and discarded from subsequent analyses (see Methods).

**Figure 2:**
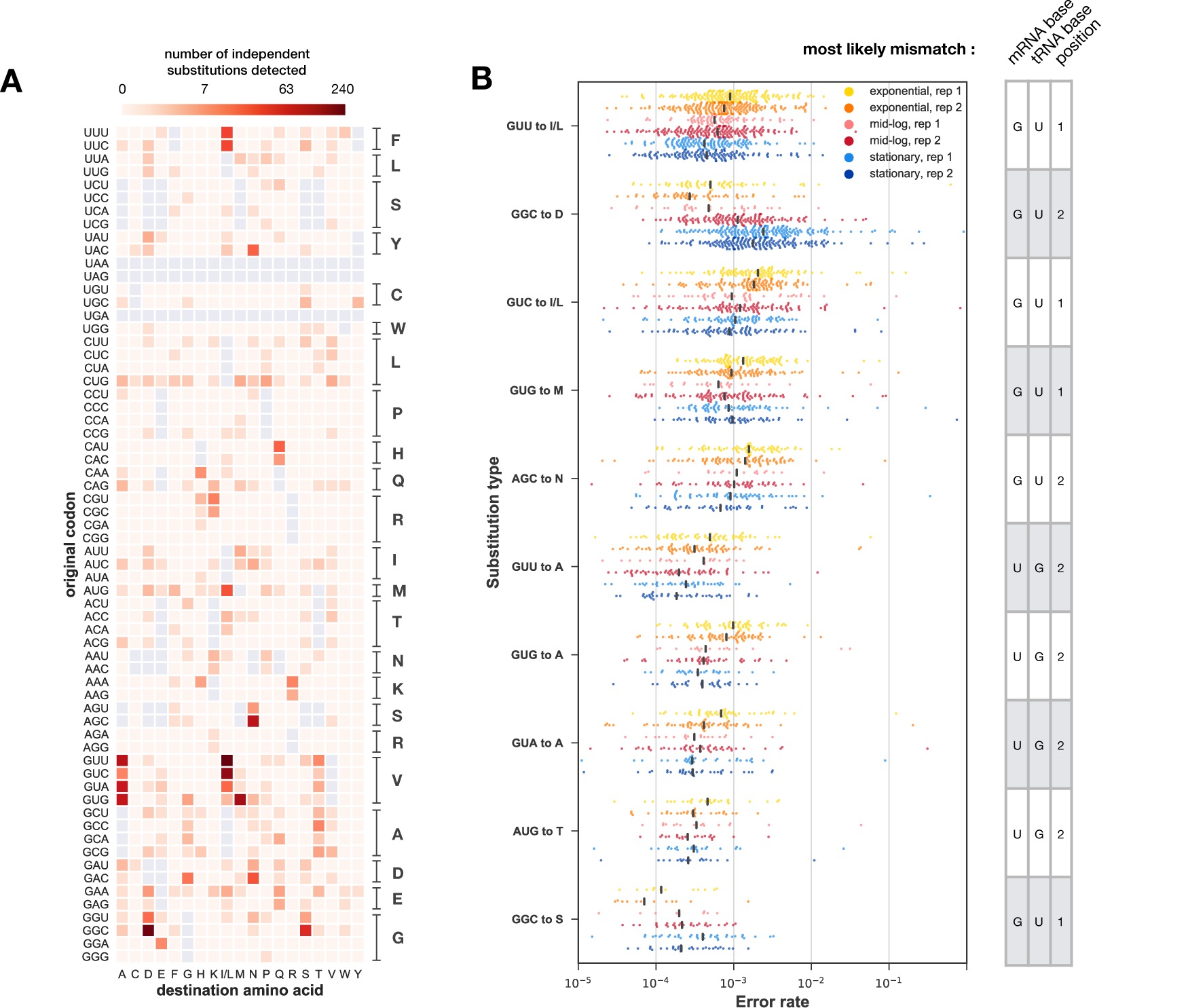
overview of the substitution profile of E. coli in MOPS complete medium. A: Matrix of identified substitutions. Each entry in the matrix represents the number of independent substitutions detected for the corresponding (original codon, destination amino acid) pair, in the MOPS-complete dataset. The logarithmic color bar highlights the dynamic range of detection. Grey squares indicate substitutions from a codon to its cognate amino acid, substitutions from stop codons, and substitutions undetectable via our method because they are indistinguishable from one of the PTMs or artifacts in the unimod.org database. Substitutions to Leu and Ile are a priori undistinguishable, and thus grouped together. B: Left panel: For each of the top 10 most frequently detected substitution types, we fetched the quantification profile of the dependent peptide and the base peptide. Each dot represents the ratio of intensities I_DP_ /I_BP_ for each of the samples, when both peaks have been detected and quantified. The black line indicates the medians of the distributions. Right panel: we inferred the most likely mismatch for each of the substitution types, using a procedure described in the Material and Methods. This allows us to guess that the V -> I/L substitutions are likely substitutions from Val to Ile, enabled by a G:U mismatch at the 1st position.

The matrix is highly structured, and some substitutions appear to be much more prevalent than others. In particular, we observe that the codon used to encode an amino acid position determines the most likely errors patterns at the corresponding site. This is nicely illustrated with substitutions from Gly to Asp and Glu. When Gly is encoded by the GGC codon, the frequent substitution destination is the near-cognate Asp (that can be encoded by the near cognate codon GAC), while encoding Gly with GGA often results in substitution of Gly by Glu (presumably due to its near cognate codon GAA). Similar cases in which different codons for the same amino acid tend to show different amino acid substitution patterns can be found in the matrix (Fig. 2A).

We calculated the observed error rate (i.e. the ratio of intensity between the dependent and base peptide) for abundantly detected substitutions. As an example, the Ser_AGC_→Asn substitution was detected among a total of 81 different peptides across the *E. coli* proteome of the non-treated samples. Figure 2B summarizes the error rate estimations in each of these substitutions – each dots in the plot corresponds to one specific SerAGC→Asn substitution on a particular genomic position, and the error rate is on the y-axis. Likewise the 10 most frequent substitutions types in the proteome are shown. The majority of the substitutions that are observable in our dataset span the error rate range around 10^−3^, with the most highly abundant substitutions types showing slightly higher error rates. Due to the MS acquisition strategy, positions that feature a low error rate are less likely to be detected, which could lead to an over-estimation of the actual error rates. We note that for most substitution types, error rates seem to consistently decline as cells progress from exponential to stationary phase – out of the 10 most prevalent substitution types, only two follow the opposite trend, and both involve the common glycine GGC codon, perhaps reflecting an intracellular shortage of glycine

### A global nucleotide mispairing pattern for translation errors

We further classified NeCE based on the type of codon-anticodon mismatch within the codon and the nucleotide types they involved, characterized by a position along the codon, a codon nucleotide and an anticodon nucleotide. We define the count density for a given mismatch type as the number of independent substitutions that can be explained by that type of mismatch, divided by the number of classes of substitutions that can be explained by the same mismatch. These count densities are presented under the form of three 4*4 “nucleotide mismatch matrices” (fig 3A), which reflect, for each of the three codon positions, the relative prevalence of the various mismatch types. Substitutions that could not be unambiguously traced back to a single mismatch type were assigned to the most likely mismatch using a greedy algorithm (see Material and Methods). The most frequently observed substitution types involve mismatches between uracil and guanine in the first or the second position of the codon. Interestingly, this rule holds mainly for G:U mismatches, where the codon base is G and the anticodon base in U; the overrepresentation of this mismatch type compared to the symmetrical U:G type could not be explained by the numerous modifications affecting the anticodon of tRNAs, as they seldom affect the anticodons’ 2^nd^ and 3^rd^ base (involved in the 1^st^ and 2^nd^ codon position mismatches)^31^

**Figure 3:**
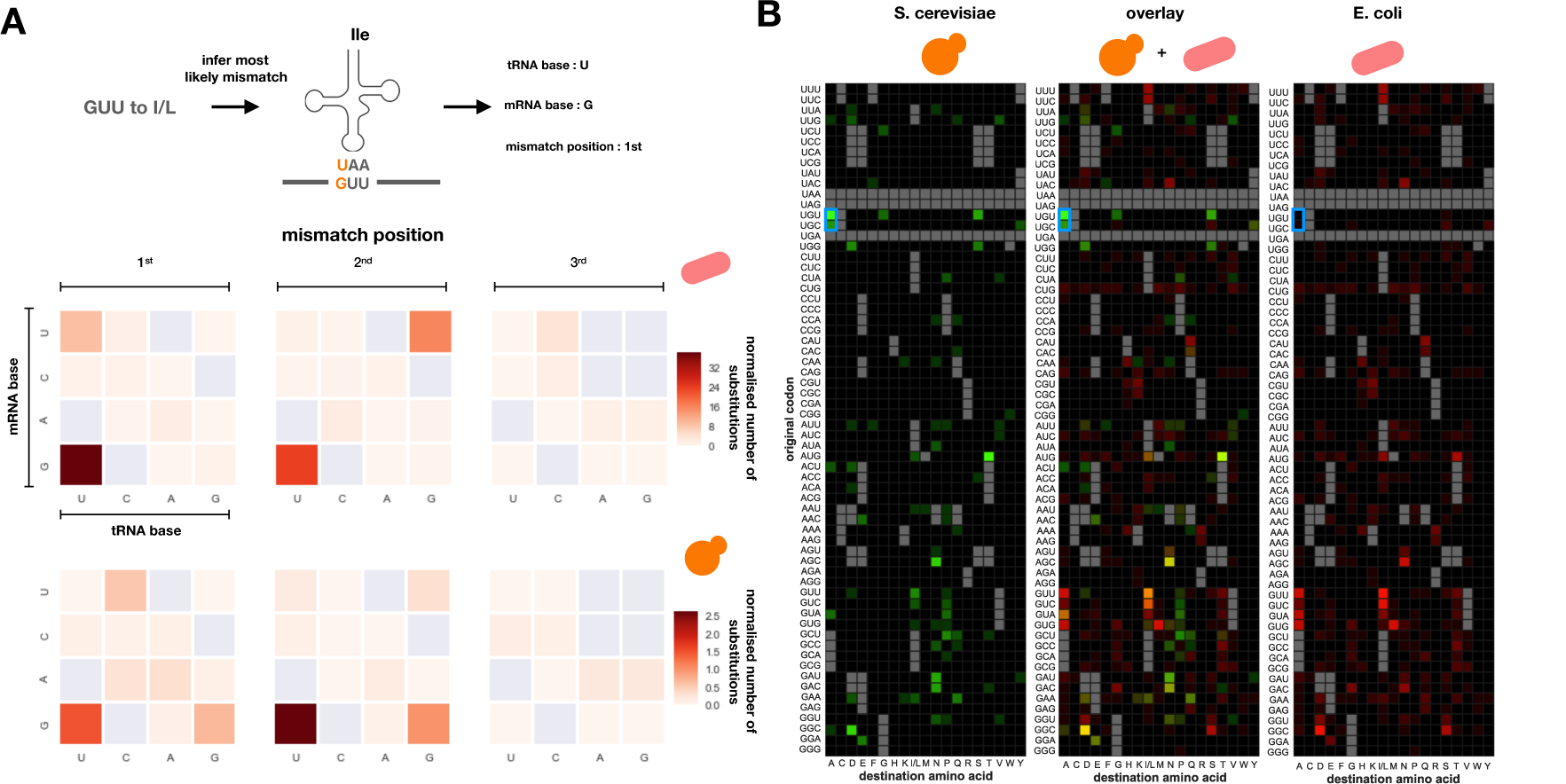
Comparing the error profiles for E. coli and S. cerevisiae reveals a shared signature of errors. A: the substitutions identifications matrices of S. cerevisiae (green channel, left) and E. coli (red channel, right) are compared and overlaid (middle). The intensity of the color is proportional to the logarithm of the number of independent identification, with one pseudo-count. Values are normalized by the highest entry in the matrix for each of the two organisms. The blue box highlights the recently described property of eukaryotic AlaRS to mischarge tRNA. B: NeCE are classified by the mismatch most likely to generate them. The shade intensity reflects the ratio of independent substitution to number of substitution types associated with the corresponding mismatch. Grey boxes are either correct base-pairings, or mismatches to which no substitutions could be unambiguously mapped. Upper panel indicates results obtained from E. coli, lower panel was generated based on S. cerevisiae data.

### *E. coli* and *S. cerevisiae* share similar error profiles

While both characterized by a mostly planktonic lifestyle and high growth rates, *E. coli* and *S. cerevisiae* have been diverging from one another for at least 2.7 billion years. Comparing the error profiles of these two organisms, thus, allows us to look at how strongly these errors are constrained, both by chemical and evolutionary necessities. We reanalyzed a previously published mass spectrometry dataset of strong anion exchange (SAX) and SCX fractionated proteome of *S. cerevisiae* grown in a single condition, a rich medium (30°C, YPD)^24^ using our pipeline.

We detected a total of 225 unique substitutions in the yeast proteome, from the two samples analyzed. Similarly to the E. coli dataset, the majority of the errors, 143, were classified as NeCE (63%). Comparing the error spectrum between the eukaryote and the prokaryote we observed a high overlap between the set of substitution types seen in the two organisms. For example, the most highly frequent substitution types (e.g. Met_AUG_ →Thr, Ser_AGC_ →Asn, Val_GUU_ →Ile) are shared between the two species. Among the 34 substitution types observed more than once in the yeast dataset, 25 had been seen in the E. coli samples. Since only 25.6% of substitution types were observed at least once in the E. coli dataset, we expected only 9 of these 34 substitution types to be detected in the E. coli data. Among the 25 substitution types also detected in E. coli, 16 were NeCE. Conversely among the remaining 9 substitution types that were not seen in the bacterial samples, only 4 were NeCE. This observation reveals a universal error pattern for mistakes that are likely to occur within the ribosome, while most NoCE likely originate from separate factors unique to each of the species. The most notable difference between the two species is in the most frequently observed substitution of Ala to Cys in yeast, which is not seen in the bacterium. A recent report^32^ revealed the basis for this observation - eukaryotic, but not prokaryotic Alanyl-tRNA synthetase (AlaRS) tend to mischarge tRNA_Cys_ with Alanine.

For the yeast data too we computed nucleotide mismatch matrices and observed that in similarity to the *E. coli* matrices they also predominantly feature G:U mismatches at the first or second positions (Fig. 4B). Observing such levels of error similarity between such loosely related organisms, exhibiting distinct codon usage biases, and relying on very different translation machineries, suggests that these errors depend on universal constraints. Whether these constraints are of a purely chemical nature, or the observed substitutions happen to be more tolerable by these organisms remains to be determined.

**Figure 4:**
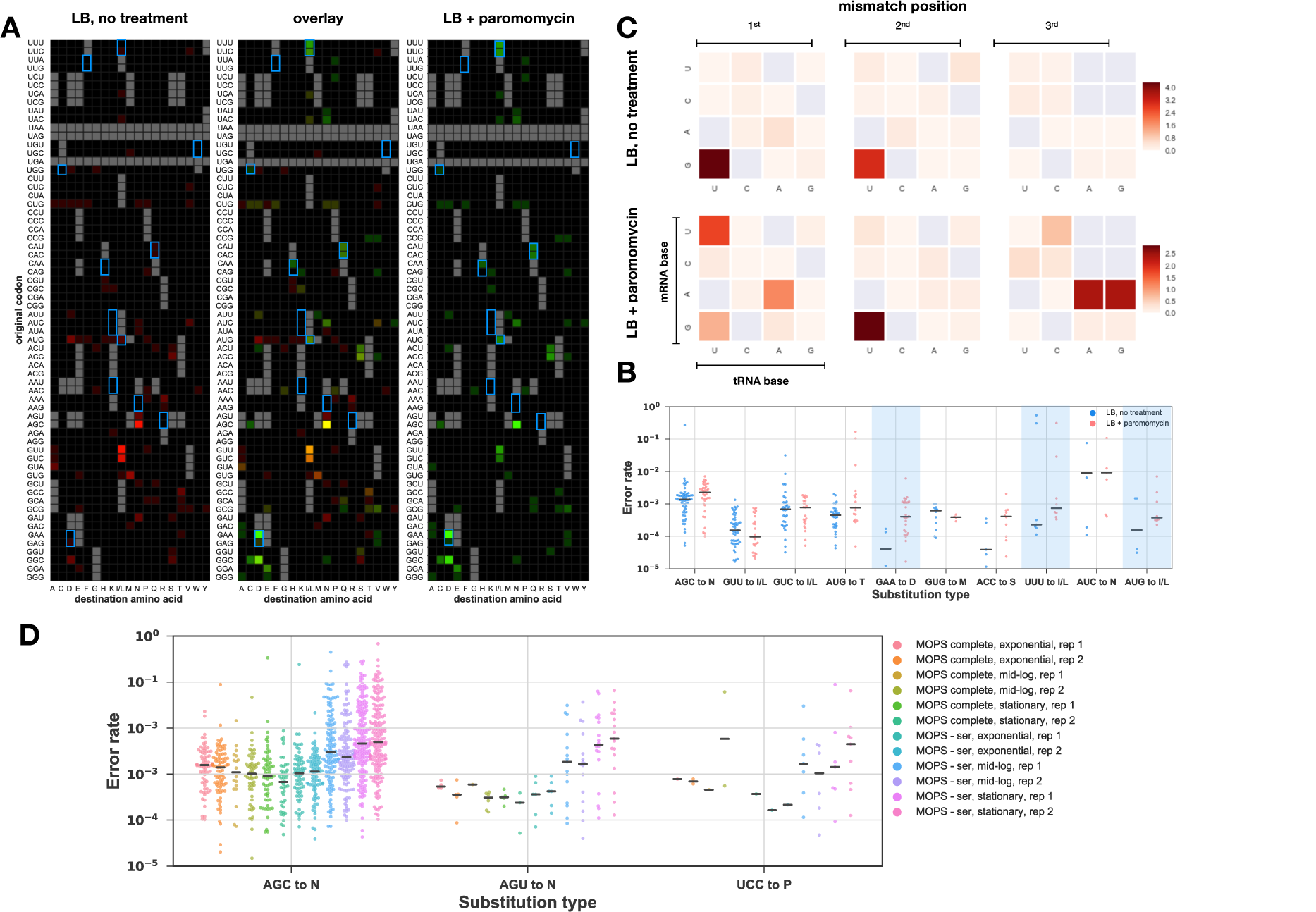
the error spectrum is affected by external perturbation. A: the substitutions identifications matrices of E. coli in LB (green channel, left), or LB supplemented with paromomycin (red channel, right), are compared and overlaid (middle). The intensity of the color is proportional to the logarithm of the number of independent identification, with one pseudo-count. Values are normalized by the highest entry in the matrix for each of the two organisms. The blue boxes highlight errors involving 3rd position mismatches. B: Quantification of the top 10 most frequent substitutions in the paromomycin dataset. Errors involving 3rd position mismatches are shaded in light blue. C: NeCE are classified using the same procedure as in Fig. 2B, for the LB samples, with or without paromomycin. D: Effect of serine starvation on errors at serine codons, for the three most frequently detected substitutions affecting serine codons.

### Aminoglycoside treatment and amino acid starvation perturb the error spectrum

We characterized the response of error patterns to external perturbations. First, we grew *E. coli* cells in LB supplemented with sub-lethal concentrations of paromomycin, an aminoglycoside antibiotic known to interfere with the ribosome’s proofreading activity^4,33^, and recorded its effect on the proteome, in comparison to that of an untreated control. For both conditions, we again inspected the codon to amino acid matrices (Fig. 4A), the error rate profiles (Fig. 4B) and the nucleotide mispairing matrices (Fig. 4C). Comparing the codon to amino acid substitution matrices between the paromomycin treated and untreated samples reveals a clear pattern – the drug increased error rates, especially at 3^rd^ codon wobble positions (Fig. 4C; greyed examples in 4B), while other mismatch positions remained relatively unaffected. The increased error rate at the 3^rd^ position can be quantified using MS1 information, as reported in Fig. 4B.

Next, we examined the impact of amino acid starvation on the cell’s error spectrum. We starved a serine auxotroph for serine, and measured its proteome. We predicted that, upon starvation to serine, we would observe elevated level of errors leading from serine to other amino acids. Indeed, the rate of Ser_AGC_→Asn markedly increased upon starvation, and intensified as the cells approached stationary phase, when the effects of starvation are supposed to be the strongest. This result indicates to a clear mechanism that accounts for mistakes in translation in which a shortage of an amino acid determines its probability to be replaced by others. While a model^34^ predicts that the UCN serine codons should be affected more strongly by starvation than the AGC and AGU codons, our method was mostly able to fetch substitutions stemming from the two latter codons. This observation does not necessarily contradict the theory, since the AGC and AGU errors are also those most seen in the MOPS complete medium, and it is possible that the UCX codons do suffer more from starvation, but remain at low levels in absolute terms. In addition, we observed that all samples in the serine starvation experiment presented a strong signal for Threonine→Serine substitutions, which did not seem to be affected by the nature of the threonine codon. Replacing threonine for serine in an environment where serine is scarce seems particularly maladaptive. However, Ling & Söll^35^ reported that E. coli’s ThrRS tended to misincorporate serine for threonine during oxidative stress. Since the starvation experiment was performed on a serine auxotroph,, we cannot exclude that the modification of the bacteria’s metabolism induced oxidative stress and therefore T→S substitutions.

### Misincorporations occur at error-tolerant and rapidly translated positions

Drummond & Wilke^18^ posited that cells, in order to avoid the fitness loss due to protein misfolding and aggregation, manage their errors by selecting error-proof codons at positions where inserting the correct amino acid is critical to a protein’s folding or function. They were able to support that theory using computational means, but as they were lacking data on actual errors in proteome they had to rely on proxies. In particular, they used evolutionary conservation as a proxy for sensitivity to phenotypic errors. They derived the identity of error prone and error proof codons from conservation data. Correspondingly, fast evolving positions within protein are predicted to be less critical for protein folding and function, and should therefore tolerate amino acid substitutions. Yet, the lack of a systematic set of translation error events within a proteome precluded so far examination of the notion that they occur in rapidly evolving sites, or in positions that minimally affect protein structure and function. A theoretical argument, kinetic proofreading^5,6^, predicts that ribosomes would be more likely to misincorporate an amino acid at sites where they translate rapidly^36^. This trade-off between speed and accuracy was demonstrated by mutants that featured modified translation speed^37^, and by in-vitro using conditions that affect ribosome velocity^38^. Even though indirect evidence suggests that, in yeast, downstream mRNA structures are used to modulate the speed of the ribosome when it decodes evolutionarily conserved sites^39^, this theory has not been experimentally tested. We therefore derived the relative speed of ribosomes at error sites via ribosome profiling^40^, and looked for direct evidence of this trade-off.

We computed normalized conservation scores for each position in the *E. coli* proteome (see Material and Methods), such that a high score indicates that an amino acid position is poorly conserved compared to other positions in the same protein. In order to account for the fact that some amino acids tend to be more conserved than other, and that some codons are over-represented at conserved positions, we devised three strategies to generate adequate negative controls (Fig. 5A). In the first control, that is least stringent, for each observed substitution, we sampled a normalized conservation score from any position in the same protein. In the second control strategy, the random re-sampling was carried not only within the same protein, but also with the additional constraint that the amino acid type in the randomly sampled position has to match the same amino acid type observed at the position at which the substitution occurred. Finally, in the most stringent of these negative controls, we performed a random re-sampling within the same protein, at sites sharing the same codon as the observed positions. We generated 1,000 such re-samplings in each of the three types of negative controls, and compared the mean of the observed distribution of scores at the observed substitution positions to those of the random control distributions to obtain empirical p-values. The mean rate of evolution at substitution sites is similar to that of random sets of positions generated though the first model, but significantly higher than that of the random sets generated with the other two (Fig. 5B). Consistent with a previous prediction^18^, controlling for the codon identity reduced the magnitude of the difference between the real error sites and randomly selected sites (Fig. 5B, “same codon” vs “same AA”), supporting the notion that evolution allows or precludes error-prone codons from sites that are correspondingly tolerant or intolerant to errors. However, codon identity did not fully account for the poor conservation at substituted sites, suggesting that other factors allow cells to locally modulate their error levels.

**Figure 5:**
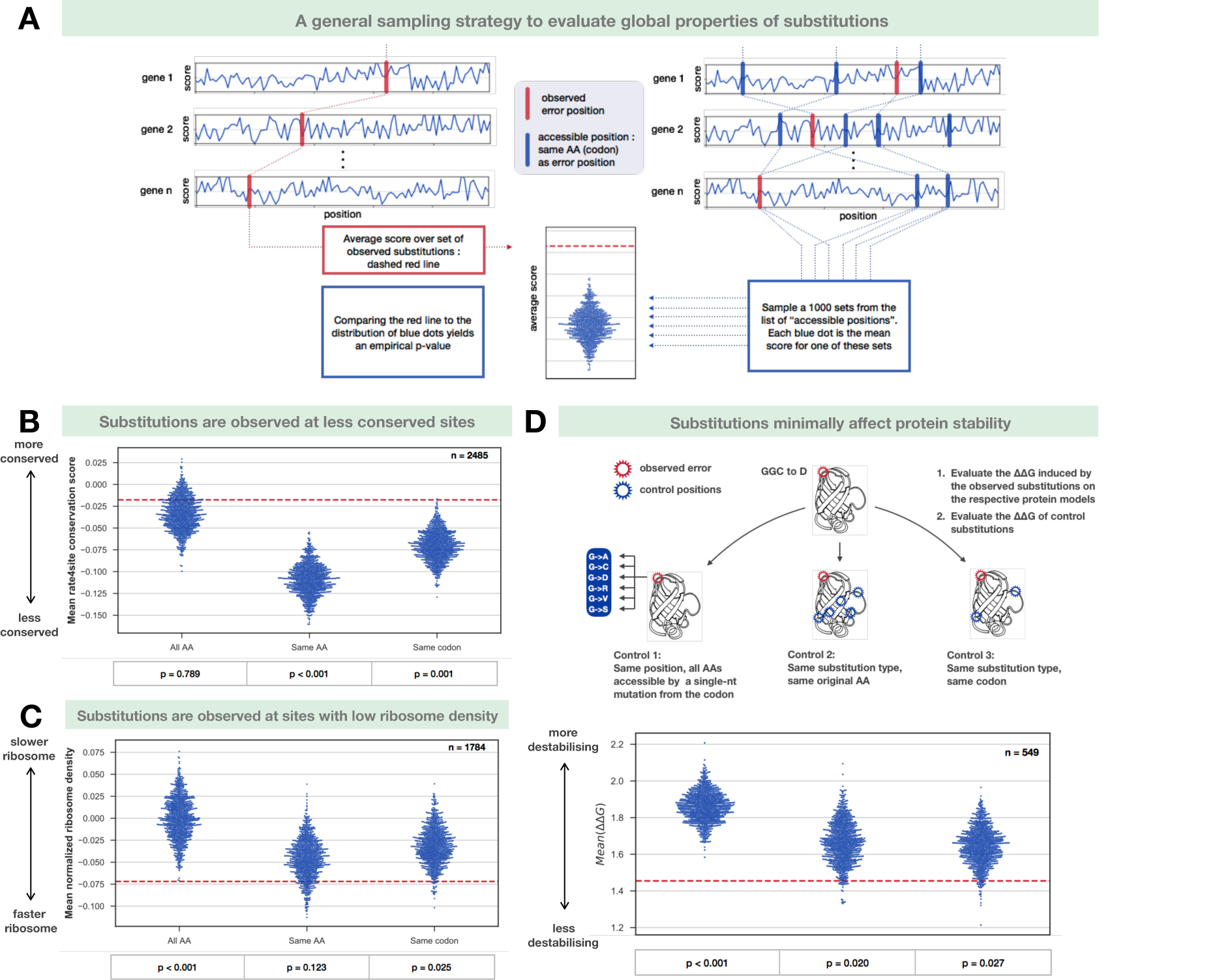
general properties of substitutions. A. The sampling strategy: In order to test if the set of detected substitutions differs from expectations in any way, we first need to account for the fact that many local properties of proteins are affected by the protein’s expression level, and so is our ability to detect substitutions from that protein. First, the local property of interest (’score’) is recorded at all the positions bearing a substitution. The average score for that set of positions is plotted as a red dashed line. To compare this average to an appropriate control, we devised three strategies to eliminate the potential contributions of protein level, amino acid identity and codon identity on the score. In each of these strategies, we draw 1000 sets of positions, and plot the average score of each of these sets as a blue dot. In the first strategy, for every bona fide distribution, we draw the score from any position within the same protein. In the second strategy, we draw the score from any position within the same protein that shares the same amino acid as the one bearing the bona fide substitution. In the third control, the codon for the sampled position also needs to be the same as the substituted codon. B. Amino acid conservation: We derived amino acid conservation scores for E. coli proteins, using the COGs database to fetch 50 homologs, MUSCLE to align them, and rate4site to estimate the evolutionary rate at each site. The resulting scores are standardized per proteins, and a high score indicates low conservation. The empirical p-values are computed by dividing the number of blue dots above the red line, divided by 1000. n indicates the number of positions considered in this analysis. C. Ribosome density: Ribosome profiling data were processed (see methods) to estimate the ribosome density at positions along the E. coli transcriptome. Since a ribosome would have to rely on local cues to modulate its speed, we do not expect misloading errors to be influenced by the ribosome density. Therefore, this analysis was restricted to NeCE. D. Effect of substitutions on protein stability: for proteins whose 3D structure is known, we evaluated the effect of NeCE on protein stability using FoldX. In control 1, we test if the observed substitutions are on average less destabilizing than those stemming from other single-nt mismatches between the codon and the anticodon, at the same position. In control 2 and 3, we test if the observed substitution type observed was less destabilizing on average at the observed position than at other positions sharing the same AA, or the same codon.

Similarly, we examined the related possibility that observed amino acids substitutions in the *E. coli* proteome tend to minimally affect the energetic folding stability of protein in which they occur. To this end, we computationally predicted the effect of each of the observed substitutions on its protein’s folding stability (ΔΔG). We compared the distribution of ΔΔG to mock distributions obtained via three control strategies, described in Fig. 5D. The first of these strategies (identity control) allows us to test if the observed NeCE are less destabilizing, on average, than other substitution types would be at the same positions. As before, we generate 1000 random sets of ΔΔG values, such that for each observed substitution we randomly sample a destination amino acid accessible via a single mutation from the error bearing codon, and compute the difference in free energy resulting from a substitution towards that amino acid. The mean ΔΔG of the set of *bona fide* substitutions, 1.454 kcal/mol, was lower than that of each of the 1000 mock sets sampled under the identity control, suggesting that error rates are tuned so that substitutions preferentially replace the original amino acid with one that would minimally affect protein folding at the same site. Our control strategy accounts for the fact that substitution types classified as NeCE tend to swap chemically similar amino acids due to the organization of the genetic code.

Next, we tested if the observed errors were preferentially mapped to protein positions were they minimally destabilized folding. We generated 1000 sets such that for each observed NeCE, we estimated the ΔΔG that would have resulted from that same substitution occurring at a randomly chosen position of the same protein, sharing either the same amino acid (“amino acid control”) or the same codon (“codon control”). The mean ΔΔG of our set of observed substitutions was lower than that of 98% of the sets generated with the amino acid control, and 97% of those generated with the codon control, suggesting that substitutions preferentially occur at protein sites where they would not disturb folding. We cannot presently exclude the alternative explanation that some substitutions destabilize protein structure, promote degradation, and are thus precluded from being sampled in our method.

The classical model of kinetic proofreading suggests that the ribosome must balance between speed and accuracy during the aa-tRNA selection step by the ribosome, and that it is more likely to misincorporate amino acids when decoding at a high rate^36^. This trade-off was experimentally observed in vitro: ribosomes tested in increasing magnesium concentrations translate their templates faster, but less accurately^41^. *In vivo*, hyper and hypo-accurate ribosomal mutants translate respectively slower and faster than the wild type ribosome^37^. Yang et al. proposed that yeast cells take advantage of mRNA structures downstream of the ribosome to tip this trade-off towards faster or more accurate decoding^39^. The speed at which ribosomes translate mRNAs can be estimated from ribosome footprints density^42^, which allowed us to test if substitutions arise preferentially at quickly translated positions.

We estimated the A-site ribosome density across the proteome of MG1655 using a published dataset acquired under similar conditions^40^, and standardized this density score per protein to control for intergenic differences in mRNA abundance and translation initiation levels (see Material & Methods). We compared the mean ribosome density at error sites to that of sets of random samples generated using the strategy described in Fig. 5A. In each of the controls, the mean ribosome density of the *bona fide* substitutions was lower than that of most of the random control samples (Fig. 5C) – error sites are less dense, and hence translated faster than expected by chance. In particular, the mean density at sites associated with a substitution was significantly (p = 0.025) lower than expected under our null model controlling for biases associated with codon identity. These results suggest that E. coli can locally tune its ribosomes to trade accuracy for speed.

## Discussion

Here we report on a new method to observe single amino acid misincorporations, which we used to detect over 3500 distinct translation errors across the proteome of *E. coli*. We take advantage of the very high accuracy of modern mass spectrometer to generate high confidence identifications. Orbitrap mass spectrometers can be tuned to detect mass differences on the order of thousandth of Daltons, during both the MS1 and MS2 acquisition phases. This accuracy in turn allows us to distinguish peptides and peptide fragments of almost identical masses, but of different atomic and isotopic compositions, and thus greatly improves the performance of database search algorithms. Our method is therefore able to distinguish amino acid substitutions from PTMs of similar masses. Despite the false discover rate (FDR) procedure applied at the end our pipeline, we cannot exclude with absolute certainty that some of the substitution types we detect correspond in fact to annotated PTMs that cannot be distinguished from amino acid substitutions. However, the retention time shifts in HPLC observed for our set of identifications correlate very well the expected retention time shifts predicted from sequence information alone, an observation that could not be explained by the identification of spurious PTMs.

One cannot guarantee *a priori* that these substitutions stem from errors in the translation machinery, because non-synonymous errors at the DNA or RNA levels could generate the same mistakes at the protein level. However, our samples originate from clonal populations, which implies that DNA mutations are unlikely to reach a detectable level in the absence of strong adaptive selection, and would be very rarely observed to occur across multiple samples. A second possibility is that we are also observing transcription errors. Yet, since we analyze samples in which the number of cells (~10^9^) is greatly superior to the inverse of the observed lower bound of transcription error rates (~10^5^), and the average number of mRNA per cell for the genes we detect is greater than one, the relative abundance of errors is expected to correspond to the transcription error rate at any examined site, and should not fluctuate from sample to sample thanks to the assumption of ergodicity. Since this estimate is two orders of magnitude lower than the average observed error rates quantified by our method, substitutions likely arise from translation errors.

Two distinct processes have been shown to generate high levels of errors in translation: aaRS can mistakenly load an amino acid to a non-cognate tRNA, and the ribosome can pair a correctly charged aa-tRNA complex to a non-cognate codon. Both processes rely on small energetic differences between correct an incorrect pairings. For the ribosome, the recognition process exploits the difference of free energy between cognate and non-cognate codon-anticodon pairs. Most aaRS also probe the nature of the anticodon of the tRNA before loading, and additionally rely on clues from the tRNA backbone to achieve a high specificity^43^. The amino acid recognition step can be challenging due to similarities between amino acid types, and a subset of these enzymes have to rely on an editing step to achieve higher specificity. Differential binding of EF-Tu to misacylated tRNAs was shown to discriminate against common aaRS mistakes^44^, and thus provides an additional layer of specificity. We argue that most of the substitutions detected in our work stem from errors in the ribosome i.e. wrong pairing between codons and anticodons. Indeed, the overwhelming majority (88%) of the substitutions could be explained by a single codon-anticodon mismatch, a fraction much higher than expected by chance due to the organization of the genetic code (30%). Additionally, we treated the cells with paromomycin, an aminoglycoside antibiotic known to perturb the accuracy of the ribosome. This drug increased the rate of several substitution types, particular those involving mismatches at the 3^rd^ codon position. These substitution types are therefore dominated by ribosome errors. However, we were able to identify several instances Cys→Ala subtitutions (NoCE) in the *S. cerevisiae* samples, consistent with a recent report that eukaryotic, but not prokaryotic AlaRS had a tendency to mischarge non-cognate cysteine tRNAs^32^.

Comparing the error spectrum of the *E. coli* and *S. cerevisiae* in untreated conditions revealed a large overlap between the set of observed substitution types, and a striking prevalence of G_codon_:U_anticodon_ mismatches at the first and second positions. Structural analysis of G:U and U:G mismatches within the ribosome revealed that they typically adopted a Watson-Crick G:C like geometry rather than the expected wobble one due to spatial constraints in the decoding center. These errors are therefore believed to originate from rare, transient geometries of nucleobases, as proposed^46^. These errors are therefore believed to originate from rare, transient geometries of nucleobases, as proposed^45^. The surprising observation that 1^st^ and 2^nd^ position G:U mismatches are typically much more prevalent than the symmetrical U:G conformation could not be explained by the many enzymes modifying the uracil on the anticodons of tRNAs, as they mostly assist the decoding of the third codon position.

The *E. coli* MOPS complete data allowed us to quantify a large number of substitutions occurring during fast growth. The mean error rate detected was on the order of 10^-3^, in the higher end of the range of previously reported estimates. Several reasons can be invoked to explain this observation. First, MS detectability is intimately linked to MS1 intensity levels: since the mass spectrometer systematically samples the most intense peptides in each scan, we are bound to preferentially detect and quantify substitutions associated to high error rates. Similarly, a peptide’s MS1 intensity depends on its abundance in the sample and on its ability to ionize well. The abundance of the correct peptide is usually much higher than that of the error-bearing one, which means that it will be sampled more often. The quantification depends on the sampling of the lower abundance, error-bearing peptide. Substitutions that increase the peptide’s ionization efficiency are therefore bound to increase its detectability, and will result in an inflated error rate. While it is generally accepted that ionization efficiency depends on a peptide’s sequence in a very non-linear fashion, we trained a linear regressor to evaluate the mean effects of amino acid composition on ionization efficiency. Our model gave satisfactory results (see appendix: Ionization Efficiency prediction), and suggests that substitutions should rarely result in a dramatic change in ionization efficiency. It remains difficult to assess to what extent the variance of the error rate distribution for each substitution type reflects biological variability or technical biases.

Comparing the median error rates of several substitutions across the different physiological states during bacterial growth revealed that they react dynamically to the changing environment: substitutions rate from valine codons tended to decrease with time, while glycine codons became more error-prone in later stages. The extent of this change might be underestimated due to the fact that we are not specifically quantifying the error rate of newly synthesized protein, but rather quantifying the errors in batch, integrating at any given time point on proteins that were synthesized till that time point that were not degraded already. Starving the cells for serine revealed a striking increase in the error rate of two substitutions involving serine codons, Ser_AGC_→Asn and Ser_AGU_→Asn. The median error rate for these codons rose to almost 10^−2^ in the stationary phase time point, with some sites reaching an error rate approaching 10^−1^. Other serine codons were also affected, but the scarcity of sampling for these rarer errors precluded a reliable quantification of the process. Theory predicts that the 4-box codons of serine (UCN) should suffer more from serine depletion than the 2-box codons (AGY) because of a differential charging of the tRNA isoacceptors^34^. Our failure to detect a large quantity of errors at TCN sites might be partially explained by the preferential usage of AGY codons in genes over-expressed during serine starvation^34^.

Translation errors have been hypothesized to affect protein evolution, and to drive the long known anti-correlation between gene expression and evolutionary rate at the protein level^18^. According to this theory, the selective pressure to prevent translation errors constrains the synonymous encoding of amino acids critical to protein folding, and organisms must select preferred, error-proof codons at positions where errors would disturb protein stability. These sensitive sites are characterized by a higher evolutionary conservation, and a slow rate of evolution. Our set of substitutions enabled us to directly test if errors are preferentially observed at fast evolving sites. Our analysis carefully controlled for the effects of protein expression level on the detectability of translation errors, the codon usage of proteins, and their evolutionary conservation. It confirmed that indeed, substitutions occur on average at less conserved sites, but also that the choice of codons could not entirely explain this effect, suggesting that other factors might affect translation accuracy in *cis*. Similarly, simulating the effects of the set of observed substitutions on protein stability revealed that they tended to be localized at sites where they minimally affected protein folding. Observed NeCE were also less destabilizing than randomly sampled NeCE at the corresponding sites, suggesting that the spectrum of ribosome errors is even more conservative than the effect of naïve single substitutions at the DNA level. Together, these results confirm that the cells encode their proteins and tune their translation machinery in ways that minimize the deleterious effects of amino acid misincorporations.

Since codon identity does not entirely account for the fact that substitutions are preferentially observed at sites where they are tolerated, we tested if the ribosome itself might modulate its accuracy locally. Several lines of evidence indicate that ribosomes optimize both speed and accuracy, and must therefore perform a trade-off between these two constraints. In particular, decreasing the ribosome’s GTP hydrolysis rate should result in an lower processing speed, but a better discrimination between cognate and non-cognate aa-tRNAs.^36^ We hypothesized that the ribosome might rely on external clues to locally slow-down in order to increase its accuracy at critical sites. Our analysis of a published ribosome profiling dataset indeed revealed a subtle but significant shift in ribosome density: the sites at which we observed substitutions were characterized by a lower ribosome density, i.e. a higher speed.

Our method provides a way to scan the proteome of organisms in various growth conditions, and can be used to unveil new types of adaptive translation^46^. In multicellular organisms, one could extend this analysis to different tissues, and diseases associated to proteostasis defects, such as Alzheimer’s^47^. Indeed, the recent report that translation error rates are inversely correlated to the maximum lifespan of rodent species indicates that maintaining high translation accuracy is critical during aging. In model organisms, it can be coupled with genetic manipulations to probe how the different components of the proteostasis network contribute to maintain accurate translation, and how errors affect the physiology and fate of proteins. This, in turn, will provide a new way to study the interplay between translation error rates and protein evolution.

## Material and Methods

### Strains and growth conditions

To generate the E. coli drugs dataset, MG1655 cells were plated on LB agar and incubated at 30°C overnight. 6 colonies of MG1655 were picked and grown until stationary phase in 3 ml LB, 30°C. All 8 cell cultures were diluted 1/100 and grown aerobically in 100 ml LB supplemented with the relevant antibiotics (see table X) in 500 ml Erlenmeyer flasks at 37°C until they reach mid-log phase (OD ≃ 0.5). For the serine starvation dataset, BW25113 (WT) and JW2880-1 (ΔserA, obtained from the Keio deletion library) cells were plated on LB agar and incubated at 37°C overnight. 2 colonies of each strain were picked and grown in 3 ml of modified MOPS rich defined medium made according to Cluzel et al recipe (SI Appendix) and incubated at 37°C until stationary phase. BW25113 and JW2880-1 cell cultures were diluted 1/1000 and grown aerobically in 220 ml of modified MOPS rich defined medium and MOPS serine starvation medium accordingly in 500 ml Erlenmeyer flasks at 37°C (mediums were made according to Cluzel et al 2012 SI Appendix).

### Proteome extraction

We adapted our proteome extraction protocol from Khan *et al.,* 2011^28^. Samples were each split into two 50 ml falcon tubes, centrifuged at 4000 rpm for 5 min, and washed twice with PBS (add 10 ml PBS, vortex, centrifuge for 5 min). Remaining PBS was vacuumed and the pellets were frozen in ethanol-dry ice. Pellets were re-suspended in 1 ml of B-PER bacterial protein extraction buffer (Thermo Fisher Scientific), pooled together, and vortexed vigorously for 1 min. The mixture was centrifuged at 13,000 rpm for 5 min. The supernatant (high solubility fraction) was collected and frozen in an ethanol-dry ice bath. The pellet was re-suspended in 2 ml of 1:10 diluted B-PER reagent. The suspension was centrifuged and washed one more time with 1:10 diluted B-PER reagent. The pellet was re-suspended in 1 ml of Inclusion Body Solubilization Reagent (Thermo Fisher Scientific). The suspension was vortexed for 1 min, shaken for 30 min, and placed in a sonic bath for 10 min at maximum intensity. Cellular debris was removed from the suspension by centrifugation at 13,000 rpm for 15 min. The supernatant was frozen in an ethanol-dry ice bath (low solubility fraction).

### SCX fractionation, HPLC and Mass Spectrometry

400μg of protein was taken for in-solution digestion and processed by Filter aided sample preparation (FASP)^48^ protocol using 30k Microcon filtration devices (Millipore). Proteins were subjected to on-filter tryptic digestion for overnight at 37°C and the peptides were fractionated using strong cation exchange (SCX) followed by desalting on C18 StageTips^49^ (3M EmporeTM, St. Paul, MN, USA). Peptides were analyzed by liquid-chromatography using the EASY-nLC1000 HPLC coupled to high-resolution mass spectrometric analysis on the Q-Exactive Plus mass spectrometer (ThermoFisher Scientific, Waltham, MA, USA). Peptides were separated on 50 cm EASY-spray columns (ThermoFisher Scientific) with a 140 min gradient of water and acetonitrile. MS acquisition was performed in a data-dependent mode with selection of the top 10 peptides from each MS spectrum for fragmentation and analysis

### Computational methods

Raw files were processed with MaxQuant v. 1.5.5.1. The list of parameters is available in the supplementary materials. High and Low solubility fractions were aligned separately. The amino acid substitutions identification procedure relies on the built-in dependent peptide algorithm of MaxQuant.

### The Dependent Peptide search

Experimental spectra are first searched using a regular database search algorithm, without any variable modification, and the significance of identifications is controlled to a 1% FDR via a target decoy procedure. Identified spectra are then turned into a spectral library, and a decoy spectral library is created by reversing the sequences of the identified spectra. For each possible pair consisting of an identified spectrum in the concatenated spectral libraries and an unidentified experimental spectrum of the same charge, and recorded in the same raw file, we apply the following steps: first we compute the mass shift Δm by subtracting the mass of the identified (unmodified) spectrum to that of the unidentified (modified) spectrum, then we simulate modified versions of the theoretical spectrum by adding *in silico* this mass shift at every position along the peptide, and finally we evaluate the match between the theoretical spectrum and the experimental spectrum using a formula similar to Andromeda’s binomial score.

For each unidentified peptide, the match with the best score is reported, the nature of the match (target or decoy) is recorded, and a target-decoy procedure^50^ is applied to keep the FDR at 1%. Peptides identified using this procedure are called Dependent Peptides (DP), whereas their unmodified counterparts are named Base Peptides (BP).

Additionally, the confidence of the mass shift’s localization is estimated using a method similar to MaxQuant/Andromeda’s PTM Score strategy, which returns the probability that the modification is harbored by any of the peptide’s amino acid.

### DP identifications filtering

The list of all known modifications was downloaded from www.unimod.org, and those marked as AA substitution, Isotopic label or Chemical derivative were excluded. Entries in this list are characterized by a monoisotopic mass shift, and a site specificity (i.e. they can only occur on a specific amino acid or on peptides’ and proteins’ termini). We removed from our analysis any DP identification that could be explained by any of the remaining modifications, using the following criteria: the recorded Δm and the known modification’s mass shift must not differ by more than 0.01 Da, and the modification must be harbored by a site consistent with the uniprot entry with a probability p ≥ 0.05. Conversely, we computed the list of all possible amino acid substitutions and their associated mass shifts. For every substitution, we only retained DP identifications such that the observed Δm and the AA substitution’s mass shift did not differ by more than 0.005 Da, and the mass shift was localized on the substitution’s original AA with p ≥ 0.95.

Among the remaining DP identifications, those such that the peptide sequence after substitution was a substring of the proteome (allowing Ile-Leu ambiguities), were also removed, to prevent pairing of dependent peptides and base peptides between paralogs. Finally, the FDR was controlled once again at 1% using the same procedure as above.

### Error rate quantification

In order to assess the error rate we quantify and compare pairs of base and dependent peptides across many samples. For each independent substitution, we fetched the quantification profile of the base peptide from MaxQuant’s peptides.txt table, and similarly fetch the dependent peptide’s quantification profile from the matchedFeatures.txt table. Whenever a peak has been detected and quantified for both the dependent and the base peptide, we estimate the translation error rate as the ratio of their MS1 intensities.

### Assignment of NeCE to their most likely nucleotide mismatch

First, for each amino acid substitution type, we fetch all the possible nucleotide mismatches that could have lead to this substitution (e.g. G:U, 1st position). Then for each nucleotide mismatch type, we count all the unique amino acid substitutions that could be unambiguously mapped to that type, and divide that number by the number of substitutions types that could be unambiguously mapped to that same nucleotide mismatch type. We call the resulting quantity the “count density”. In subsequent iterations of the algorithm, we repeat the previous assignment procedure, but in cases where a substitution can be mapped to more than one nucleotide mismatch type, we break the tie based on the count density of the corresponding nucleotide mismatch types, and repeat until convergence.

### Evolutionary rates computation

For each of the proteins associated to a substitution in the MOPS dataset, we fetched a list of orthologous protein sequences from the COG database^51^, excluding partial matches (membership class = 3). Proteins whose list of orthologs contained less than 50 sequences were excluded from this analysis. For the remaining proteins, we randomly selected 50 sequences from the list, and created evolutionary alignments using MUSCLE^52^. The alignments were then used to compute normalized evolutionary rates per site with the rate4site program^53^. The mean evolutionary rate of sites associated with detected substitutions was compared to that of a 1000 randomly sampled positions, using the strategy described in Fig. 5A

### Effect of substitutions on protein stability

For each of the proteins associated to a substitution in the MOPS dataset, we attempted to fetch the best 3D structure for its biological assembly in the PDB database to estimate the effect of our substitutions on protein stability using the FoldX software^54^. We excluded membrane proteins, whose stability is poorly modeled by FoldX, and excluded ribosomal protein because the ribosome is too big to be modeled entirely. We restricted our analysis to WT proteins from E. coli, excluding structures determined from orthologs. Among the remaining structures, we selected those with the lowest R-free score.

These structures were first “repaired” using the repairPDB command. We then evaluated the effect of a set of amino acid substitutions comprising the detected substitutions and the controls described in Fig. 5D on protein stability (ΔΔG), using the PositionScan command. Finally the mean(ΔΔG) of our set of substitutions was compared to the mean(ΔΔG) of 1000 randomly sampled substitutions, using the strategy described in Fig. 5D.

### Ribosome density computation

Ribosome profiling data for the MOPS complete experiments were downloaded from Woolstenhulme *et al.*, 2015^40^ (GSM1572266, GSM1572267). Reads were aligned using the 3’ mapping method described in the corresponding article, and shifted by 12 nt to obtain the density at the A-site. Read counts from both replicates were summed to obtain more robust estimates, and 20 codons were excluded from both the 3’ and the 5’ ends to avoid known biases. For the remaining positions, we applied the transformation x : log_2_(x + 1) to stabilize the variance, and standardized the resulting score to obtain the normalized read density (NRD), so that the mean of the NRD per protein is 0 and its standard deviation is 1. The mean(NRD) of the set of observed substitutions was then compared to that of 1000 randomly sampled substitutions, using the strategy described in Fig. 5A.

